# Rapid Liquid Chromatography-Mass Spectrometry (rLC-MS) for Deep Metabolomics Analysis of Population Scale Studies

**DOI:** 10.1101/2025.08.10.669154

**Authors:** JD Watrous, S Tiwari, T Long, A Large, K Lagerborg, K Gallagher, K Dao, S Park, D Aguirre, C Lamoureux, J Cho, R Nilsson, J Usuka, M Jain

**Affiliations:** Sapient Bioanalytics, LLC; 10421 Wateridge Circle, Suite 100; San Diego, CA 92121

## Abstract

Mass spectrometry (MS)-based metabolomics is a key technology for the interrogation of exogenous and endogenous small molecule mediators that influence human health and disease. To date, however, low throughput of MS systems have largely precluded large-scale metabolomics studies of human populations, limiting power to discover physiological roles of metabolites. Here, we introduce a fully automated rapid liquid chromatography-mass spectrometry (rLC-MS) system coupled to an AI-enabled computational pipeline that enables high-throughput, reproducible, non-targeted metabolite measurements across tens of thousands of samples. This system captures thousands of polar, amphipathic and nonpolar (lipid) metabolites in a human plasma sample in 53 seconds of analytical time, enabling analysis of greater than 1,000 samples per day per instrument. To demonstrate the discovery power of the rLC-MS platform, a subset of samples from Sapient’s DynamiQ™ biorepository – comprised of 62,039 total plasma samples collected longitudinally from 11,045 individuals – were selected for deep analysis by rLC-MS to capture a rich, dynamic landscape of chemical variation that reflects both physiological processes and environmental influences. 26,042 plasma samples with matched real-world data (RWD) were chosen for the study, representing 6,935 individuals with diverse demographic backgrounds and disease profiles. Unbiased exploratory analysis revealed human metabotypes that correlate with heterogenous disease phenotypes, including key sub-populations of cardiometabolic and other human diseases. Moreover, a metabolic aging clock machine learning model trained on healthy individuals in this dataset accurately predicted accelerated aging in various chronic diseases, with dynamic reversal of metabolic aging following definitive therapy. These data demonstrate that the rLC-MS platform enables prediction of clinically relevant physiological states from plasma metabolomics at scale in human populations.

## Introduction

Advances in mass spectrometry (MS) resolution and sensitivity have enabled increasingly comprehensive views into the diverse molecular and biochemical landscape that influence human health and disease ^1,2^. Whereas traditional ‘targeted’ MS based metabolomics approaches focus on a pre-determined set of well-characterized molecules, including amino acids, bile acids, oxylipins, structural lipids, as well as drug metabolites, discovery or ‘non-targeted’ high resolution MS-based technologies are now being leveraged to provide a more comprehensive view of human health via the ‘undiscovered’ metabolome, measuring thousands of small molecule metabolites in a single biospecimen ^3,4^. Non-targeted MS has the potential to provide critical insights into mechanisms by post-genomics, dynamic influences, including diet, lifestyle, environmental exposures, toxicants, microbes, as well as host genotype and biochemistry influence on human biology, disease pathobiology and drug response ^1,2,5^. In addition to providing mechanistic insight into human biology, small molecule metabolites may also serve as powerful diagnostic and prognostic predictors of human disease, owing in part to their diverse exogenous and cellular origins, rapid translocation into central circulation, and dynamic nature ^5,6^.

To date, analytical throughput has largely precluded application of non-targeted MS to diverse, population-scale datasets, limiting statistical power, robust cross validation and overall discovery potential. While in-line / off-line solid phase extraction systems or direct MS without chromatographic separation (‘flow injection’) allow for very fast sample-to-sample MS injection times ^7,8^, such approaches limit quantitation of molecules due to space charging, complex matrix interactions, and limited overall system sensitivity. Similarly, alternative analytical approaches such as NMR, while enabling rapid and highly reproducible metabolomics measures, are often restricted to measure of the most abundant molecules in complex biosamples ^2^. For metabolomics analysis of human biosamples, liquid chromatography coupled to mass spectrometry (LC-MS) has typically been employed to enable stationary retention and separation of diverse small molecules prior to MS analysis ^1^. While effective, standard chromatography methods often take 20-60 minutes per sample and are usually optimized to measure a specific class of compounds (such as amino acids, lipids, fatty acids, or carbohydrates), requiring multiple chromatographic separations and mass analysis, with concatenation of spectral data post-acquisition ^1^.

Herein we describe a non-targeted rapid liquid chromatography (rLC) system coupled to high resolution, ion-mobility quadrupole time-of-flight mass spectrometry (rLC-MS). This system provides rapid capture and separation of diverse metabolite chemistries — from polar to amphipathic and nonpolar (lipid) metabolites — in a single injection while overcoming the bottlenecks in current LC-MS methodologies, enabling robust and reproducible measurement of thousands of small molecules in human plasma in less than one-minute analytical cycle time per sample. In a population scale survey of 26,042 plasma samples acquired across time from 6,935 diverse individuals, rLC-MS data provides a unique view into the landscape of human small molecules, revealing both endogenous physiology and environmental influences. We find that circulating metabolites hold close association with key human health and disease phenotypes, including the ability to subset complex disease phenotypes by distinct metabotypes. Moreover, a metabolic aging clock trained on this data enables prediction and readout of complex, dynamic phenotypes and biological processes, including biological aging, disease onset and therapeutic response.

## Results

### Human plasma metabolomics at scale

To enable measurement of the plasma metabolome across populations at a scale that enables robust human discovery, an ideal analytical system must have high throughput, capture a broad spectrum of compounds, yield reproducible results over long time periods, and be robust against variation in sample handling. Current LC-MS methods are constrained by several bottlenecks that limit overall throughput, including serial, independent chromatography gradients, mobile phase flow rates, sample acquisition, and data transfer between the LC-MS and computing systems. To overcome these limitations, we developed a rapid liquid chromatography (rLC) system leveraging stationary phase column chromatography with multiple retention mechanisms acting in a ‘mix mode’ fashion, coupled to a high resolution, ion-mobility (IMS) quadrupole time-of-flight (QToF) mass spectrometer (Fig. 1a). The rLC chromatography system retains a broad range of chemistries in human plasma, ranging from hydrophilic sugars, amines and organic acids to amphipathic and neutral lipids (Fig. 1b), within an analytical cycle time of 53 seconds. To evaluate the sensitivity and dynamic range of the rLC-MS system, we introduced eight labeled standards spanning a variety of chemical classes (Supplementary Table 1) into a plasma sample at 0.1 to 1 times the reported physiological concentrations ^9^. All standards demonstrated linearity over this concentration range, down to high picomolar concentrations (Fig 1c, Suppl Fig 1a,b). Moreover, in serial dilution of a pooled human plasma sample, 80% of all non-targeted rLC-MS features demonstrated linearity (Pearson *r* > 0.8) over a 10-fold sample dilution range (Suppl Fig 1c,d), with linearity independent of mass or chemical characteristics, including retention time (Suppl Fig 1e,f). rLC-MS measurements were also minimally affected by hemolysis and lipemia (data not shown). Collectively, these data suggest that rLC-MS enables measures of plasma small molecules with high fidelity across a broad concentration range.

**Figure 1.**
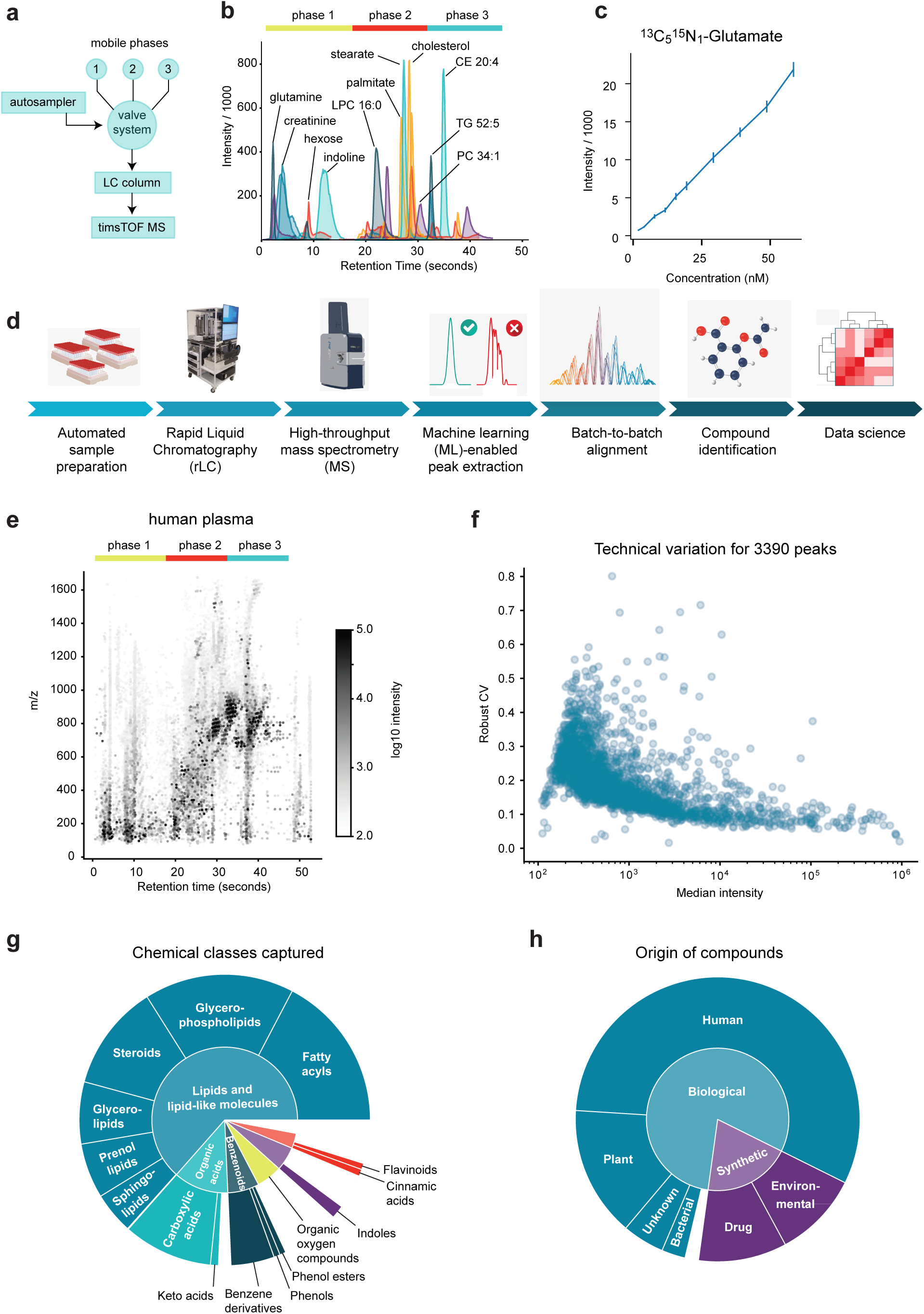
Large-scale human plasma metabolomics with the rLC-MS platform. **a**, Schematic of the rLC-MS system. Three mobile phases are sequentially introduced via a valve system into the LC column, coupled to a high-resolution timsTOF mass spectrometer. **b**, Extracted ion chromatograms for selected metabolites, with rLC chromatography phases indicated. **c**, Example internal standard dilution curve for ^13^C_5_^15^N_1_ - glutamate in positive ionization mode. Error bars denote standard deviation over n = 10 replicates. **d**, Schematic of the full rLC-MS pipeline. **e**, Retention time, mass/charge ratio (m/z) and intensity distribution of 30,139 peaks extracted by the rLC-MS pipeline from a single human plasma sample. **f**, Technical variation of rLC-MS peak intensity for 3,390 peaks extracted from a pooled plasma sample injected 891 times across 6 batches. **g**, Breakdown of 1,424 annotated features by classification using the ChemOnt ontology. **h**, Distribution of identified metabolites by origin.

To enable high throughput discovery, we integrated the rLC-MS into a complete system platform consisting of an automated sample handling system and a downstream data processing pipeline leveraging machine learning (ML)-enabled automated peak detection ^10^, batch-to-batch alignment and compound identification (Fig 1d). This ML system routinely captures ∼30,000 rLC-MS spectral features (extracted ion chromatograms) in typical human plasma samples (Fig 1e) with a false positive rate of 2% relative to manual ‘expert’ data evaluation. In repeated preparation and rLC-MS analysis of a single human plasma sample over several months by several different operators, we observed a median coefficient of variation (CV) in peak height of 17%, with more intensive peaks demonstrating less variability (Fig 1f), indicating good reproducibility. To identify metabolites on the rLC-MS platform, we assembled and characterized an extensive in-house library of 11,837 synthesized, chemically diverse standards. Of these, we obtained high-confidence metabolite identifications for 1,424 compounds in human plasma when matching by retention time, MS1, MS2, and collision cross-section (CCS) information (Supplementary Table 2). These compounds span a variety of chemical classes (Fig 1g), including endogenous human metabolites, bacterial metabolites, natural products and drugs (Fig 1h). These results demonstrate the ability of rLC-MS to capture diverse chemistries originating from both endogenous and exogenous sources in human plasma.

### The chemical landscape of a human population

To explore the potential of the rLC-MS platform for metabolic profiling of a population scale human cohort, we selected a well-characterized subset of individuals from Sapient’s DynamiQ^TM^ biorepository that is comprised of 62,039 samples from 11,045 individuals (Table 1). We analyzed 26,042 plasma samples acquired across multiple timepoints from 6,935 individuals with diverse demographic backgrounds (Supplementary Table 3). Biosamples were matched with comprehensive, longitudinal real world data (RWD) and deep clinical phenotyping data, including demographic information, diagnoses, medications, lab measures, procedures, clinical outcomes, and health survey data. Within the subset of 26,042 plasma samples, 3.7 samples on average were collected per individual over a three-year time frame and analyzed using the rLC-MS platform in 6 independent sample batches. Machine learning-based peak detection followed by filtering of background peaks present in blanks and removal of likely in-source fragments, adducts and mass isotopomers yielded in a comprehensive set of 15,439 rLC-MS features representing distinct metabolites and lipids. For these features, biological variation among individuals was markedly higher than technical variation in measurement (Fig 2a), indicating substantial power to discern biological effects. Importantly, unbiased clustering of samples did not group samples by batch or run order (Suppl Fig 2a,b), indicating that rLC-MS data is reproducible and comparable across batches.

**Figure 2.**
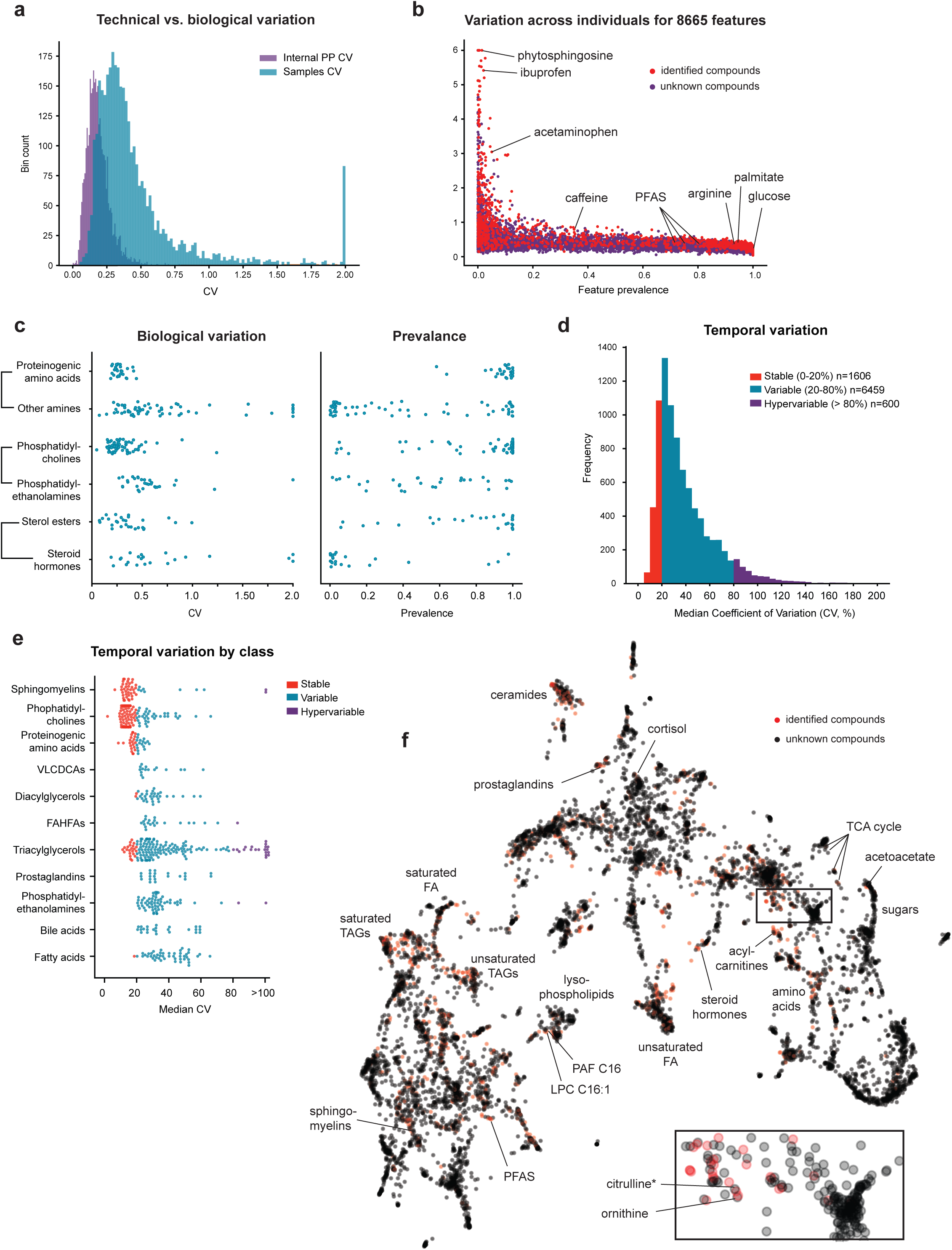
Chemical landscape of a human population. **a**, Distributions of variance (CV) across 26,042 human plasma samples and across 891 pooled plasma (PP) injections. **b**, Individual variation quantified as CV and prevalence (fraction of samples where detected) for all 8,665 common features across individuals in the cohort. For indicated metabolites, see text. **c**, CV and prevalence for selected classes of metabolites. Brackets indicate related metabolite classes. **d**, Histogram of metabolite temporal variation, defined for each feature as the median of CV within each individual over serial sampling across all individuals. **e**, Breakdown of temporal variation by metabolite classes. **f**, UMap projection of pairwise correlations across the 8,665 features. Identified features highlighted in red; selected metabolites and metabolite groups indicated. Bottom right, zoom-in of area indicated by rectangle.

**Table 1.** Demographics and selected lab measures in DynamiQ.

| Variable | No. Individuals | No. Samples | Mean (±SD) | Fraction |
| --- | --- | --- | --- | --- |
| Total | 11,045 | 62,039 |  |  |
| Age | 11,040 | 61,055 | 55.3 (±15.3) | |
| Gender |  |  |  |  |
| Female | 6,483 |  |  | 58.8% |
| Male | 4,544 |  |  | 41.2% |
| Race/Ethnicity |  |  |  |  |
| White | 5,525 |  |  | 50.1% |
| Hispanic | 2,161 |  |  | 19.6% |
| Asian | 1,988 |  |  | 18.0% |
| Black | 833 |  |  | 7.6% |
| Other | 533 |  |  | 4.8% |
| Variable | No. Individuals | No. Measurements | Mean (±SD) | |
| Body Mass Index | 10,691 | 134,361 | 27.3 (±6.0) | |
| Systolic Blood Pressure | 9,559 | 195,672 | 124.8 (±18.3) | |
| Diastolic Blood Pressure | 9,557 | 194,791 | 74.3 (±11.8) | |
| Hemoglobin A1c | 7,216 | 41,212 | 6.0 (±1.2) | |
| LDL cholesterol | 7,519 | 54,083 | 101.4 (±38.0) | |
| Triglycerides | 7,595 | 55,402 | 131.6 (±162.6) | |
| eGFR | 9,004 | 211,225 | 57.1 (±35.7) | |

To examine the spectrum of metabolic variation between individuals, we quantified both prevalence, defined as the fraction of individuals in which an rLC-MS feature was reliably detected (Suppl Fig 2c,d), and individual variation (CV) across the 6,935 individuals (Fig 2b, Suppl Fig 2c). Rare (low prevalence) but highly variable metabolites included exogenous molecules such as pharmaceutical agents, as well as diet-derived molecules such as plant products. In contrast, endogenous central human metabolites such as glucose, amino acids and major fatty acids were prevalent and varied less across individuals. Polyfluorinated compounds that derive from exposure to industrial chemicals and persist in body tissues for long periods of time (“forever molecules”) were also remarkably common in this population (Fig 2b), suggesting widespread exposure. Proteinogenic amino acids were remarkably constant compared to other amines (Fig 2c), suggesting homeostatic regulation of these central metabolites. Similarly, structural lipids such as phosphatidylcholines showed little variation while phosphatidylethanolamines were more dynamic, and steroid hormones were found to be more rare and variable than other sterol molecules (Fig 2c).

To assess the temporal dynamics of metabolites in human plasma, we analyzed plasma samples from 1,126 individuals with at least 5 independent blood samples collected over greater than a year, computed CV across time points for each metabolite within each individual, and took the median CV across these 1,126 individuals as a measure of temporal variation. Overall, the distribution of metabolite temporal variation within individuals (Fig. 2d) was similar to variation across individuals. Among metabolites that were stable over time (CV < 20%) were structural lipids such as phosphatidylcholines and sphingomyelins as well as the proteinogenic amino acids (Fig 2e). The majority of metabolites investigated varied to some degree over time (CV 20%–80%), including bioactive and signaling lipids such as very-long-chain dicarboxylic acids (VLDCAs), diacylglycerols and prostaglandins (Fig 2e). Interestingly, triacylglycerols were spread among stable, variable, and hypervariable (CV > 80%) temporal patterns, suggesting different sources or specialized biological functions.

To investigate functional relationships between metabolites, pairwise correlations of all features were computed across all samples and this correlation structure projected into 2D using the UMap algorithm (Fig 2f), reasoning that functionally related metabolites should co-occur across individuals and across time. This analysis revealed a rich landscape of interrelated metabolites, crossing chemical families. For example, immunologically active but structurally unrelated metabolites such as prostaglandins and corticosteroids clustered together, suggesting that abundance patterns of these metabolites reflect immunological state of individuals. Saturated triacylglycerides (TAGs) clustered near saturated fatty acids and separate from unsaturated TAGs, possibly reflecting variations in dietary fats. We also found direct biochemical relationships to be reflected in feature clustering, with for example platelet-activating factor (PAF) located next to its precursor LPC 16:1. Among polar metabolites, the central organic acids of the TCA cycle formed a tight cluster located near the ketone body acetoacetate and sugars/aminosugars. Such co-clustering of related metabolites also helps identify metabolites: for example, one rLC-MS feature putatively annotated as citrulline or argininic acid clustered nearby ornithine, the immediate precursor of citrulline in the urea cycle, suggesting that this feature is in fact citrulline. Thus, leveraging feature correlation structure in large human cohorts can both identify metabolites and recover physiologically relevant groups of metabolites.

### Data-driven metabotypes predict disease states

A long-standing question in human metabolism is whether there exist subpopulations of individuals with characteristic metabolic phenotypes, or “metabotypes”, and whether these metabotypes may reveal altered risk for development of common diseases ^11,12^. To investigate this question, we first selected a small subset of rLC-MS features that were prevalent in the population scale cohort and yet demonstrated high biological variation (CV > 0.5) across individuals, resulting in 385 features. These included fatty acids and triglycerides of different chain length and composition, as well as gut microbiota-derived metabolites (Fig 3a). To define metabotypes in a data-driven manner, we performed unsupervised clustering of all 6,935 individuals based on correlations between these feature vectors (Suppl Fig 3). This grouped individuals into six clusters with striking and consistent differences across the metabolite groups (Fig 3a). From here on, we refer to these clusters as metabotypes. Metabotype 1 exhibited high levels of short TAGs as well as fatty acids, while metabotypes 2 and 3 showed either high TAG levels and low free fatty acids, or vice versa, suggesting differences in lipid metabolism between these subpopulations. Metabotype 4 stood out with high abundance of a distinct cluster of features that contained several gut bacteria-derived metabolites, while metabotype 5 was characterized by high abundance of long TAGs but low short TAGs and free fatty acids. Metabotypes were found to be largely independent of common demographic features, and for instance did not differ markedly in age distribution, except for metabotype 6 that contained a younger population (Fig 3b). Moreover, other metabotypes with distinct metabolite patterns shared demographic measures, with metabotypes 1 and 4 both associated with high BMI (Fig 3c), despite their molecular differences. These data indicate that large-scale rLC-MS metabolomics data has the power to discern individuals with specific metabolic states across populations.

**Figure 3.**
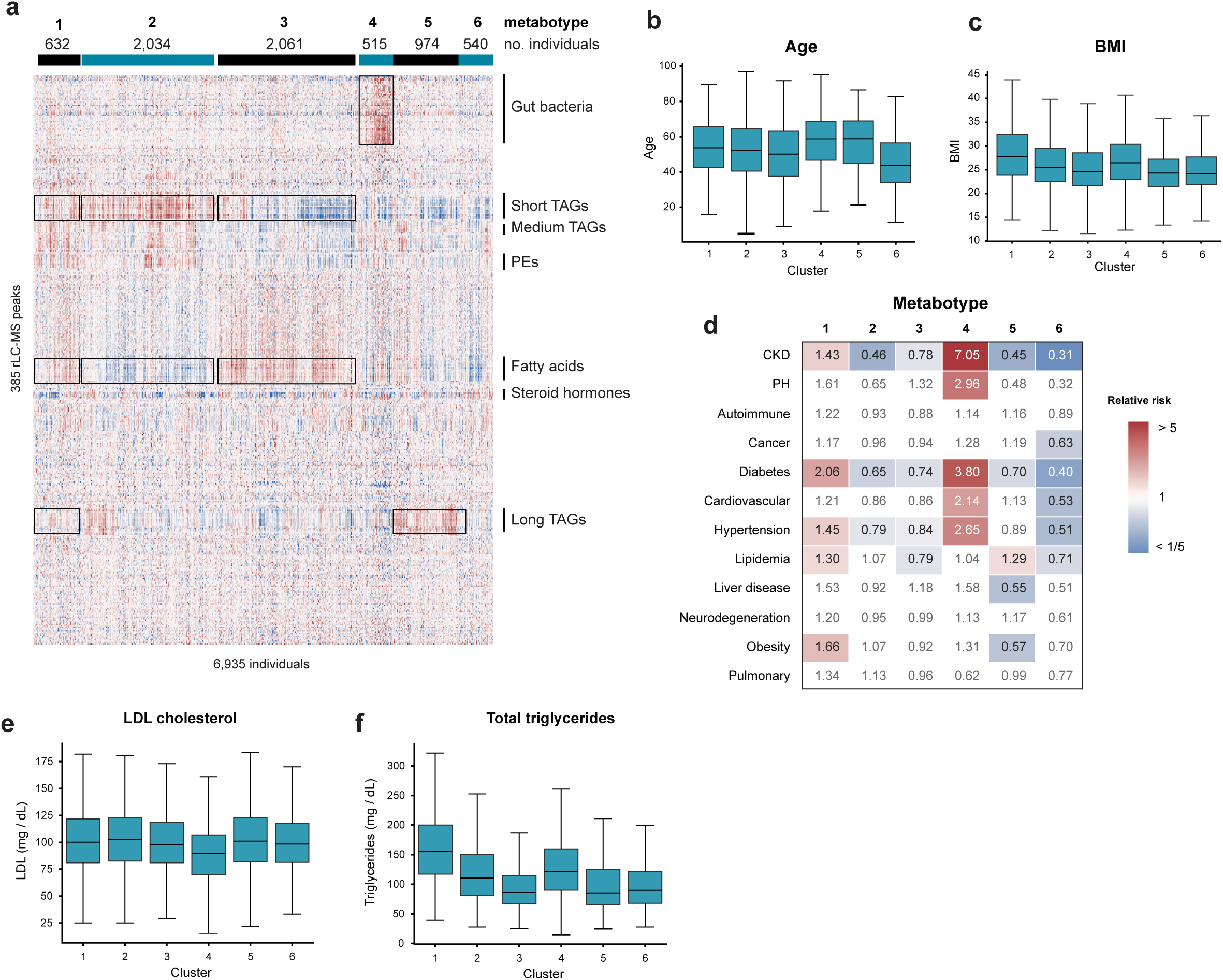
Unsupervised discovery of metabotypes in a human population. **a**, Clustered heatmap of 385 prevalent and highly variable rLC-MS features across 6,935 individuals. Mean values of all samples for each individual are shown. Top, metabotypes obtained from cluster analysis, excluding 179 individuals not assigned to any cluster. Right, selected metabolite groups (see text). **b-c**, Distribution of age (b) and body mass index (BMI; c) for individuals in each metabotype. **d**, Heatmap of relative risk for indicated prevalent disorders for each metabotype. Non-significant associations are shown in gray text. **e-f**, Distribution of LDL cholesterol (e) and total triglycerides (f) for individuals in each metabotype.

To investigate whether these metabotypes might be related to common human disorders (Table 2 and Supplementary Table 4), we determined the relative risk of prevalent disease for each metabotype and found they were strongly associated with cardiometabolic diseases (Fig 3d), but not with autoimmune or neurodegenerative diseases. In particular, metabotype 4 showed high risk of cardiovascular disorders, diabetes, and in particular chronic kidney disease, but was not significantly associated with traditional cardiovascular risk factors, including lipidemia or obesity. Interestingly, LDL-cholesterol was significantly lower in metabotype 4 (Fig 3e), despite the high prevalence of cardiovascular disease in this group. In contrast, metabotype 1 was associated with obesity, lipidemia and diabetes, but showed no significant increase in cardiovascular risk. Metabotype 5, characterized by abundant long TAGs, was also associated with lipidemia, and yet showed lower risk of obesity, diabetes and liver disease. Interestingly, total triglycerides were in the normal range for metabotype 5, but elevated in metabotypes 1 and 4 (Fig 3f). These data demonstrate that our database of human plasma metabolomes contains clearly distinguishable subgroups, which can predict metabolic disorders in an unsupervised fashion from metabolomics data, independent of traditional disease risk factors.

**Table 2.** Selected diseases in DynamiQ.

| **Common disease** | **No. Individuals** | **No. Samples** |
| --- | --- | --- |
| Cancer | 1,649 | 11,570 |
| Diabetes | 2,550 | 16,906 |
| Obesity | 2,360 | 13,338 |
| Lipidemia | 4,408 | 29,863 |
| Liver disease | 928 | 5,738 |
| Hypertension | 4,306 | 28,682 |
| Cardiovascular | 3,281 | 23,975 |
| CKD | 2,583 | 17,774 |
| Pulmonary disease | 2,061 | 13,609 |
| Autoimmune | 844 | 5,718 |
| Neurodegenerative | 724 | 4,609 |
| **Rare Disease** |  |  |
| Cystic fibrosis | 7 | 82 |
| Amyloidosis | 111 | 916 |
| Pulmonary hypertension | 564 | 4716 |
| Amyotrophic lateral sclerosis | 6 | 50 |

### Metabolic aging clock predicts accelerated aging

Given the dynamic nature of plasma metabolites, which derive from endogenous, dietary and environmental sources (Fig 1h) as well as integrate information on biological processes across multiple body tissues, we next asked if population-scale rLC-MS metabolomics data can predict complex physiological traits. As a case study, we considered biological age, a data-driven measure that captures individual differences in rate of aging. While biological age was initially defined based on epigenetic markers ^13^, predictors of aging from plasma proteins ^14^ or metabolites ^15–17^ might better capture dynamic changes in aging due to disorders. To predict biological age, we trained a machine learning model on 1,640 samples from 887 healthy individuals of ages from 23 to 82 years old having no diagnosed chronic diseases and BMI between 18.5 and 30, for whom biological age should closely align with chronological age. To obtain an interpretable model that can be implemented using targeted methods, we used the Stabl framework ^18^ to select a set of 30 rLC-MS features that together produced an accurate multivariate model, referred to as the metabolic aging clock (see Methods). On a fully independent lockbox validation set of 758 samples from 300 individuals, this model predicted individuals’ age with a median absolute error of 8.0 years (Fig 4a). Among predictive features negatively associated with age were several sex hormones (Fig 4b), which are known to decline with age. Positively associated metabolites included cystine, the oxidized form of cysteine, in line with reports that thiol oxidation increases in aging ^19^; glucuronate, which is involved in detoxification of xenobiotics; and phenylacetylglutamine, a marker of nitrogen surplus which has recently been implicated in senescence during the aging process ^20^. For several other metabolic markers in the age prediction model, the biological roles remain to be explored.

**Figure 4.**
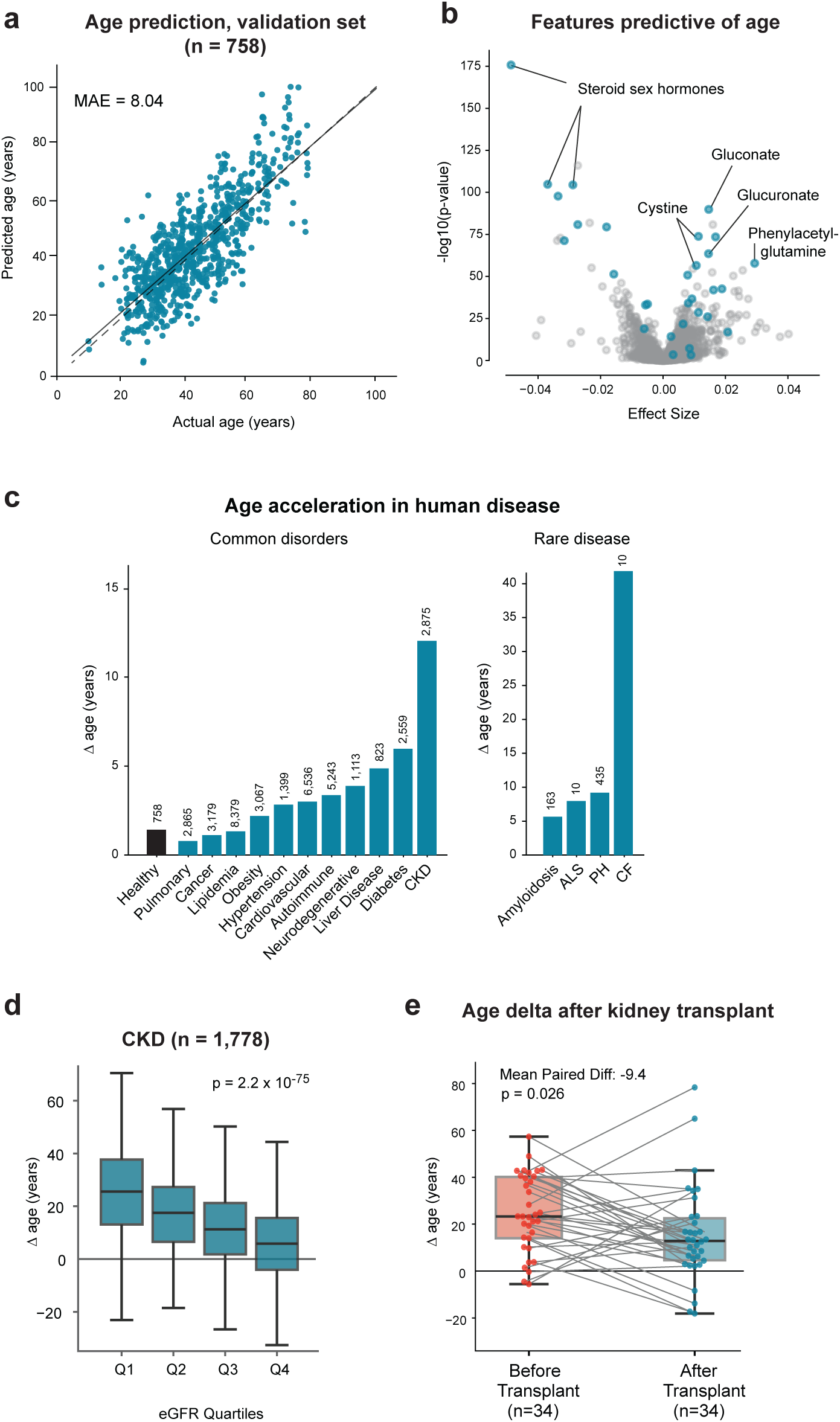
**a**, Performance of the metabolic aging clock model on the lockbox validation set of 758 samples from 300 healthy individuals. Solid line indicates fitted linear regression model; dashed line, identity function. MAE, median absolute error. **b**, Effect size (regression coefficient) vs. p-value for linear regression association between each feature and chronological age, adjusted for gender and BMI, in the training/testing set. Features selected for inclusion in the metabolic aging clock model are highlighted; identified metabolites are indicated. **c**, Age acceleration (Δage), defined as difference between predicted and actual age, of individuals with indicated chronic disorders. Median values per group are graphed; number of samples per group are indicated. CKD: chronic kidney disease. ALS: amyotrophic lateral sclerosis. PH: pulmonary hypertension, CF: cystic fibrosis. **d**, Age acceleration *vs*. quartiles of estimated glomerular filtration rate (eGFR) for individuals diagnosed with CKD. **e**, Age acceleration in 34 individuals before and after kidney transplant. Gray lines indicate samples from the same individual.

Chronological age generally cannot be perfectly predicted from biomedical data since individuals of the same age may differ widely in physiology and biological age. Because common human diseases tend to accelerate aging (shortening lifespan), we asked if age predictions by our model would be affected by disease. In a set of 4,000 individuals not seen by the model during training, we estimated an acceleration in biological age (Δage) of several years for individuals diagnosed with common chronic disorders (Fig 4c, Suppl Fig 4a, Table 2 and Supplementary Table 4). For example, obesity was associated with a median Δage of 3.7 years; diabetes 7.4 years; and chronic kidney disease as much as 13.0 years, consistent with the reported reduction in overall lifespan associated with each of these disease states. Moreover, within individuals diagnosed with kidney disease, Δage was inversely correlated with estimated glomerular filtration rate (eGFR; Fig 4d), a continuous measure of renal dysfunction, indicating that within this disease state, more severe disease was related to more accelerated aging. Similarly, among individuals with hyperlipidemia, higher plasma triglycerides were associated with higher Δage (Suppl Fig 4b). This suggests that biological age predicted by our metabolic aging clock reflects disease severity in a more fine-grained manner than traditional diagnosis. In contrast, diseases generally localized to single organs with less effect on systemic physiology exhibited a more minimal effect on accelerated aging. For instance, non-metastatic cancer increased median Δage by only 2.4 years. Other diseases that primarily manifest in single organs, including rare debilitating diseases such as amyotrophic lateral sclerosis (ALS) or cystic fibrosis, resulted in substantial accelerated aging, suggesting widespread effects on whole body function (Fig 4c). These data suggest that the metabolic aging clock captures intrinsic aging factors that are affected by systemic diseases. We next asked whether our model would reflect a ‘reversal’ of accelerated aging after therapeutic treatment of disease and thereby inform on the overall treatment response. To this end, we applied the metabolic aging clock to a set of individuals with end-stage renal disease that underwent kidney transplantation. Remarkably, the biological age predicted from plasma samples of these individuals decreased markedly after transplantation (Fig 4e), suggesting that pro-aging factors affecting systemic aging were normalized with definitive treatment. This decrease was not due to drift of Δage with time, since Δage was stable over a one-year period among healthy individuals (Suppl. Fig 4c). Taken together, these data demonstrate that large-scale plasma profiling of human populations with the rLC-MS method can predict biomedically relevant physiological states.

## Discussion

Our results demonstrate that the rLC-MS method captures a wide range of human metabolites with a throughput of ∼1,000 samples per day per rLC-MS system, enabling deep analysis of human plasma metabolomes at scale. The ability to map metabolomes across tens of thousands of samples for large cohorts brings new opportunities, as large sample size is often critical for discovery in human populations. We envision that rLC-MS metabolomics will substantially increase the power to identify small molecule biomarkers and discover novel disease mechanisms. Importantly, we find that data generated from the rLC platform is reproducible across batches of samples run at different times and by different operators. This is critical for the development of both biomarkers and machine learning models, for example to predict risk for disease or response to drugs, as any such predictive model must naturally be applicable to future batches of data. In addition, the availability of large collections of human metabolomes allows exploring the correlation of rLC-MS features to identify novel metabolites by similarity to known ones, and to elucidate the role of metabolites of interest in physiological processes.

Our exploratory analysis of a large human cohort indicates that there exist well-delineated metabotypes in human populations which can be discovered from rLC-MS metabolomics data alone by a simple, unsupervised clustering analysis. These metabotypes appear to reflect innate metabolic differences, such as metabotypes 2 and 3 (Fig 3) that show markedly different plasma triglyceride and free fatty acid content, suggesting that metabotype 2 individuals preferentially transport fatty acyls as lipoprotein-carried triglycerides, while metabotype 3 individuals have higher plasma free fatty acids, possibly due to differences in lipolysis and esterification. Metabotypes were also differently associated with prevalent disorders, further indicating that they are physiologically distinct. Our analysis focused on a small fraction of the most prevalent and variable rLC-MS features that happen to include major blood lipids such as triglycerides and fatty acids, which likely explains why these particular metabotypes show associations with metabolic disorders but not with autoimmune or neurodegenerative disease. It must be emphasized that associations between metabotypes and human disorders do not imply causality: for example, it is quite possible that high abundance of the gut microbiota cluster of metabolites in metabotype 4 (Fig 3) is a consequence of chronic kidney disease among these individuals rather than a cause. Also, our analysis did not adjust for medications such as statins, which could skew the results if drug use differs between groups. Nevertheless, it is clear that there exist metabolically defined subgroups of individuals that cross conventional diagnosis lines, such as metabotypes 1 and 4 which are both associated with diabetes and hypertension, but where metabotype 1 exhibits obesity and lipidemia but not heart disease while metabotype 4 shows the opposite pattern. These data suggest that plasma metabolomics profiles can be used to develop molecular diagnosis tools that more accurately reflect an individual’s physiological state, and which could inform on personalized treatment.

Plasma metabolomics data is an excellent resource for building artificial intelligence models to predict various aspects of an individual’s physiological state, as plasma metabolite content is highly dynamic and reflects both diet, lifestyle, environmental exposures and endogenous metabolic processes. The high throughput of the rLC-MS method is crucial in this application, as machine learning methods require large numbers of training examples to generate accurate models. While most rLC-MS features are unidentified, as is common in metabolomics data, the entirety of the data can nevertheless be used as a “fingerprints” of physiological processes. As an example, we demonstrate the ability to predict biological age from rLC-MS data using an interpretable linear model. Our metabolic aging clock, trained on rLC-MS metabolite profiles from 887 healthy individuals, successfully predicted accelerated aging in chronic human disorders as well as reversal of aging in individuals with kidney disease following transplantation. This example demonstrates that it is indeed feasible to train predictive models of complex physiological states such as aging, even from limited rLC-MS data sets. We envision that, as larger cohorts of rLC-MS data become available, increasingly sophisticated machine learning models can be developed for key outcomes such as onset of disease and response to drug treatments.

## Methods

### Sample Preparation

For all plasma studies, de-identified human blood samples were collected under standard protocols, plasma was isolated and frozen, and transported on dry ice. Samples were thawed, aliquoted into Matrix tubes (Thermo) and then stored at –80 °C until sample processing. A VisionMate 2D barcode reader (Thermo) was used to record and confirm sample locations. Plasma samples were thawed over 10 minutes using a custom-made Rapid Plate Thaw system and placed on an orbital shaker at 550 rpm at 4 °C for 10 minutes. An Agilent Bravo liquid handling system was used to transfer 16 μL of sample to a shallow-well 96-well plate containing 200 μL of extraction solution consisting of 65:35 Acetonitrile:Methanol with 0.5% formic acid and containing the following internal standards: ^13^C_5_^15^N_1_ -Glutamate (Sigma-Aldrich), 16:0-d31-18:1 PC (Avanti Polar Lipids), 14:0-16:1-14:0 d5 TG (Avanti Polar Lipids), Arachidonic Acid-d11 (Avanti Polar Lipids), and CUDA (Cayman Chemicals); see also Supplementary Table 1. Samples were shaken at 550 rpm at 4 °C for 10 minutes followed by centrifugation at 6000g at 4 °C for 10 minutes. Supernatants were then transferred to a 384-well polypropylene plate containing either 80:20 Water:Methanol with 0.05% formic acid and 1mM ammonium formate or 60:40 Water:Methanol with 0.1% acetic acid for positive and negative mode analysis, respectively. Additional isotopically labelled standards were incorporated into the dilution step, namely Phenylalanine-d5 (Cayman Chemicals), and MAPCHO-12-d38 (Avanti Polar Lipids). The diluted extracts were then centrifuged at 6000g at 4 °C for 10 minutes to remove any remaining precipitate and transferred to a foil-sealed 384-well plate. Four racks of 96 tubes were processed concurrently into one 384-well plate.

For each rack of 96 tubes, three pooled plasma samples were included for quality control purposes together with 93 samples, these were prepared alongside samples using the same liquid handler and procedure described above. For each 384-well plate an additional external bracket (commercial pooled plasma) was prepared by hand using the same procedure described above and analyzed at the beginning of each run.

### rLC-MS System

Mobile phase was introduced into a custom packed silica-based mixed mode column using three independent, continuous, high flow pumps using a high fidelity nanovalve switch adjacent to the column and optimized to limit dwell volume and flow delay, resulting in a tri-phasic step gradient that gradually changed ionic strength, pH, solvent viscosity, and flow rate over time. Samples were introduced into the column using a flow-through, vacuum-assisted aspirator with sample-to-column time limited to 0.3 seconds with injection stacking to limit dwell time. The rLC instrument was coupled to a high-resolution ion-mobility quadrupole time-of-flight (QToF) tims-TOF Pro 2 mass spectrometer (Bruker) scanning over an m/z range of 50-1800, with paired systems operating in positive and negative electrospray ionization (ESI) modes. The mass spectrometer parameters were set as follows: dry gas temp of 350 °C, dry gas flow rate of 8(+) / 6(-) L/min, nebulizer gas of 50 psi, sheath gas temp of 150(+) / 250(-) °C, sheath gas flow rate of 3 L/min, source voltage of (+)3500 / (-) 3000 V, End Plate voltage of 2000(+) / 1500(-) V, mass range set to 50 – 1700 m/z, and data were collected at 9 spectra/s.

To limit time required for coupling the rLC-MS instrument to storage computing and file indexing, data was acquired as a continuous single file across 384 samples, with data segmentation performed using sensor logs post-capture. Collectively, these modifications enabled analysis of biological samples in a rapid, automated manner with a 53 second analytical cycle time per sample. All samples were run in parallel on two rLC-MS instruments with dedicated configurations for either positive or negative mode sample analysis.

### rLC-MS Quality Control

Quality control (QC) of spectral data was performed using a panel of isotopically labeled internal standards (Supplementary Table 1) as well as pooled plasma samples to monitor fluctuations in extraction efficiency, instrument sensitivity, matrix artifact and mass accuracy. Mass calibration was performed prior to each 384-well plate run and used to assess mass accuracy, mass resolution, detector sensitivity, and instrument cleanliness. For each 384-well plate run, isotopically labeled internal standards were added to each sample at the first sample preparation step to monitor matrix effects. Bulk pre-aliquoted pooled plasma was placed in three wells of each 96-well plate and prepared identically to samples (internal bracket QC samples). Bulk pre-aliquoted commercial pooled plasma was prepared external to the 96-well plate by hand (external bracket QC sample), and a preparation blank was prepared during each 384-well plate to assess background. Internal and external bracket QC samples were used to determine if any drift observed in plasma samples originated from the automation in the sample preparation process. The internal bracket QC samples and the internal standards added to samples were used to determine if any source of drift observed in samples was inherent to the samples or a result of sample preparation. Overall coefficient of variance (CV) for the internal standards were below 20% with the exception of 16:0-d31-18:1 PC which was below 25%.

### Dynamic Range Experiments

To investigate linearity and dynamic range, eight labeled standards spanning a variety of chemical classes (Supplementary Table 1) were introduced into pooled human plasma, spanning a range from 0.1x to 3x physiological concentrations, for a total dynamic range of ∼2.5 orders of magnitude. For each concentration, 10 replicate samples were prepared and analyzed. For linearity of plasma metabolites, a pooled plasma sample was extracted using varying amounts of extraction solution to achieve a range from 0.1 to 3 times normal concentrations.

### Spectral Data Handling

Chromatographic drift was assessed and corrected for based on common “landmark” features observed in all samples ^21^. Metabolite features were then detected and extracted using customized machine learning software ^10^. The core of this pipeline leverages a UNet model that segments chromatograms into regions corresponding to distinct chromatographic peaks, generating binary masks that delineate peak boundaries. The UNet model was trained on a curated dataset of 20,000 manually “expert” labeled rLC-MS spectra from human plasma samples, encompassing a diverse range of chemical classes and signal intensities. False positive rates were assessed by comparing ML-detected peaks against manual expert evaluation on a subset of 4,001 spectra, achieving classification area under the curve (AUC) for peak detection above 0.993. Operating at true positive rate of 96.6% at bin level, the corresponding false positive rate was 2.00%.

Post-segmentation, features were extracted as the highest intensity of the signal in the window selected by the segmentation algorithm, followed by batch median normalization to correct for plate-to-plate batch related effects in feature intensities. This step yielded a preliminary set of 104,442 rLC-MS features detected in at least one of the 26,042 samples. To ensure quality, features were filtered by requiring a minimum intensity of 500 counts and 3x the median intensity observed in blank samples in at least one sample, reducing the set to 63,542 features.

To further identify and filter adducts, isotopomers, and fragments in the rLC-MS dataset, a pipeline that integrates peak mass spectrometry metadata with peak intensity data was developed and utilized. Predefined mass differences for common adducts (e.g., [M+H]^+^, [M+Na]^+^), isotopes (e.g., ^13^C shifts), and fragments (e.g., loss of H₂O, CO₂) were compiled into a reference table and filtered by polarity where applicable. Peak metadata, including m/z and retention time was cross-referenced with this table to identify potential parent-child relationships, using tolerances of ±10 ppm for m/z and ±0.05 minutes for retention time. Potentially related peaks were then confirmed by computing Pearson correlations between their intensity values across samples, flagging those with a correlation coefficient ≥ 0.7 and a p-value ≤ 0.05.

Naturally occurring isotopomers were identified by estimating the number of carbon atoms in each parent peak using an empirical linear model based on m/z, followed by predicting expected peak height ratios for ^13^C isotopes based on natural abundance (1.109%). Observed ratios (child peak intensity divided by parent peak intensity) were compared to these predictions, retaining isotope annotations only when within an acceptable range. The final annotated dataset flagged each peak with associated adducts, isotopomers, or fragments, grouping them into parent and child relationships. These filtering steps resulted in a reduced set of 15,439 rLC-MS features.

To focus on features common among batches, a set of 8,665 features were retained by keeping features with intensity higher than 500 in at least 5 samples in one batch and in at least three batches.

For the metabolic aging clock, missing values for each feature were imputed from a uniform distribution ranging from half of the lowest to the lowest intensity observed for that feature across all samples. For unbiased clustering, missing values for each feature were resampled from all non-missing values for that feature. The former method better models missingness due to low abundance, while the latter preserves each feature’s marginal distribution and ensures batch-to-batch reproducibility.

### Standard Library and Metabolite Identification

A standard library was constructed by curating 11,837 small molecules selected from the Human Metabolome Database (HMDB), PubChem, and LipidMaps to achieve broad coverage of human endogenous metabolites, FDA-approved drugs, food additives, supplements, environmental toxicants, pollutants, and other environmental exposures. Standards were analyzed using the rLC-MS system in both positive and negative ionization modes. Sample preparation and dilutions were performed on a Bravo automated liquid handling system (Agilent Technologies), with final compound concentrations optimized to 0.2–1 µM. Quality control measures included daily calibration and equilibration of the rLC-MS system and inclusion of five blank and ten quality control injections per batch prior to analysis of standards. Each 384-well plate run included 12 internal QC injections to monitor run quality and concluded with one blank injection. This process generated 299,539 extracted ion chromatograms, including adducts. MS2 spectra were acquired in both polarities, resulting in a library of 22,476 spectra. MS2 data acquisition was performed using the TIMS Control software (Bruker) with methods tailored to compound m/z ranges: for compounds with m/z > 200 Da, parallel accumulation-serial fragmentation (PASEF) mode was used, while for compounds with m/z ≤ 200 Da, data-dependent acquisition (DDA) mode was employed. MS2 spectra were collected using a single collision energy that was stepped up as the m/z increased, with a 1 Da precursor ion isolation window. In addition, MS2 spectra were acquired from a set of human plasma samples in parallel reaction monitoring (PRM) mode using custom methods for large-scale MS2 collection.

Spectral data were curated using an in-house neural network model trained on manually annotated standards, with extracted ion chromatograms (EICs) as the primary feature and human-labeled categories of ‘real signal’ or ‘noise’. The training dataset was augmented by shifting retention times in both directions to prevent the model from bias in selecting features in a specific retention time region. Model performance was evaluated with an area under the curve (AUC) threshold of >0.90, ensuring only high-quality signals were used for downstream annotations.

Features in human plasma data matching library standards were manually reviewed based on feature shape consistency, retention time consistency, and high signal-to-noise ratio. MS2 spectra from human plasma were matched against the in-house MS2 library using cosine similarity, with manual verification of major fragment matches to confirm identifications.

### Study Cohort

Sapient’s DynamiQ biorepository is comprised of 62,039 longitudinal plasma samples obtained from 11,045 community dwelling individuals recruited from a medical center in an urban area. The study was conducted under Institutional Review Board ethical approval. All participants provided written informed consent at the time of study enrollment. For this study, a subset of 26,042 well-characterized plasma samples were selected from 6,935 individuals to represent diverse demographic backgrounds and disease profiles. Plasma samples were assayed using Sapient’s rLC-MS platform, as described above, and all data are deposited in Sapient’s DynamiQ multi-omics database. Metabolomics measures were integrated across timepoints to provide dynamic views of small molecule biomarker patterns associated with each individual’s changing health and disease trajectories. Demographic information such as age, gender, race, and ethnicity were collected at enrollment using standardized questionnaires and linked to each sample in the database. Clinical information was obtained from electronic health records and integrated with the molecular measures to enable deep phenotyping approaches and stratification of individuals into subgroups. Data handling and storage procedures were designed to protect the confidentiality and privacy of participant data and comply with all relevant regulatory rules and guidelines.

### Metabolite Temporal Variation

To acquire a reliable measure of temporal variation in circulating metabolites, a subset of 11,397 samples serially collected over a year time period from 1,126 individuals were analyzed. The coefficient of variation (CV) for each feature for each individual over time was calculated and the median of CV for each feature across individuals was used as a measure of temporal variation. Features with a median CV below 20% were labeled as “stable”; features with median CV between 20% and 80% were labeled as “variable”; and features with median CV above 80% were labeled as “hypervariable”. Enrichment analysis was performed as follows: for each compound class and for each label, a 2×2 contingency table (member/not a member of the compound class vs. with a given label/without a given label) was created. One-sided Fisher’s exact tests were then performed on each contingency table. Compound class with at least 4 members and p-value < 0.05 was considered “enriched” for a particular temporal variation label.

### Metabolite Feature Clustering

For metabolite feature clustering, imputed data was normalized to the median of each feature and log2-transformed. The resulting data was projected to 2 dimensions using the UMap algorithm as implemented in the umap-learn python package v 0.5.7, with the Pearson correlation metric and 15 nearest neighbors.

### Metabotype Clustering and Disease Enrichment

Metabotypes were defined using unsupervised clustering of samples as follows. To focus on features that were likely to be informative for clustering analysis, a set of 385 features having prevalence > 0.2 and CV > 0.5 was selected. Imputed data was collapsed to a single vector per individual by averaging across all samples derived from each individual, normalized to the median of each feature, log2-transformed, and clipped to the range [-3, 3] to prevent single outliers dominating the distance measure. This resulted in a 385 x 6,935 matrix of relative feature intensities for each individual. Hierarchical clustering was applied to rows and columns of this matrix using the Pearson correlation distance measure and average linkage. Metabotypes were defined by cutting the clustering tree for individuals to obtain 15 distinct clusters and discarding any cluster containing less than 50 individuals, resulting in 6 clusters as metabotypes.

Relative risk for each disease and for each metabotype were calculated by considering the individuals belonging to the metabotype as in the study group, and all other individuals as in the reference group. Significance of relative risk was calculated using Fisher’s exact test, and p-values < 0.05 after Bonferroni correction for the total number of tests were considered significant. Disease groups were defined as in Table 2.

### Metabolic Aging Clock

To build a sub cohort comprised of healthy individuals for training/testing and validation, two steps were taken. First, individuals diagnosed with diseases listed in Table 2, in the past or during follow-up, were excluded. Second, individuals with body mass index (BMI) under 18.5 and over 30 were excluded. This resulted in 1449 individuals. 300 individuals with 758 samples were reserved as a holdout lockbox set for validation. From the remaining pool, individuals were down-sampled to have equal number of samples within each age bracket across 20-30, 30-40, 40-50, 50-60, and 60+ years of age. This resulted in a training/testing set of 887 individuals with 1,640 samples. Training/testing was split with 70% and 30% and stratified by age bracket as well as individuals to ensure balance between sets.

To focus on common features across all batches, 3,768 features were selected as the initial set for building the metabolic aging clock model. For each feature, a linear regression was fitted between age and the feature intensity, with covariates of gender and BMI in the training/testing dataset. False discovery rate (FDR) correction was applied using the Benjamini-Hochberg method, with annotated and unannotated features corrected separately. This resulted in 1,458 features with FDR < 0.05 that were passed to the Stabl framework, using elastic net as a base estimator. Stabl selected a set of 32 robust features, which were then fed into three different machine learning models in sklearn: lasso regression, random forest regression, and xgboost regression. Optuna was used to select the machine-learning model with hyperparameter optimization that best reduces the mean absolute error (MAE) between actual and predicted age. To prevent overfitting, a penalty term was added as 50% of the difference between the train MAE and the test MAE. This resulted in a lasso regression model with 30 features. This model was calibrated to reduce the bias at the extremes of the age distribution by fitting a linear regression between the predicted and actual ages from the training set, and using the intercept and slope to correct for the bias.

Lastly, the final model, designated as the metabolic aging clock, was applied to the healthy lockbox validation set to evaluate the model performance, and to individuals with chronic diseases to estimate accelerated aging. For each disease listed in Supplementary Table 4, individuals with blood samples collected between 90 days prior to diagnosis and 5 years after diagnosis were selected. Metabolic age was predicted and age delta – the difference between metabolic and chronological age – was calculated for each sample from these individuals with chronic diseases. To assess whether accelerated aging was observed for a given chronic disease, a Student’s t-test was performed on age delta to compare with the healthy lockbox validation set.

To investigate the association between age acceleration and the common clinical biomarkers among individuals diagnosed with common chronic disorders, BMI, blood pressure, and other lab measurements were selected when the measurement date was within three months of sample collection date as summarized in Supplementary Table 3. Association analyses were performed with linear regressions with outliers removed.

---

**Supplementary Figure 1.**
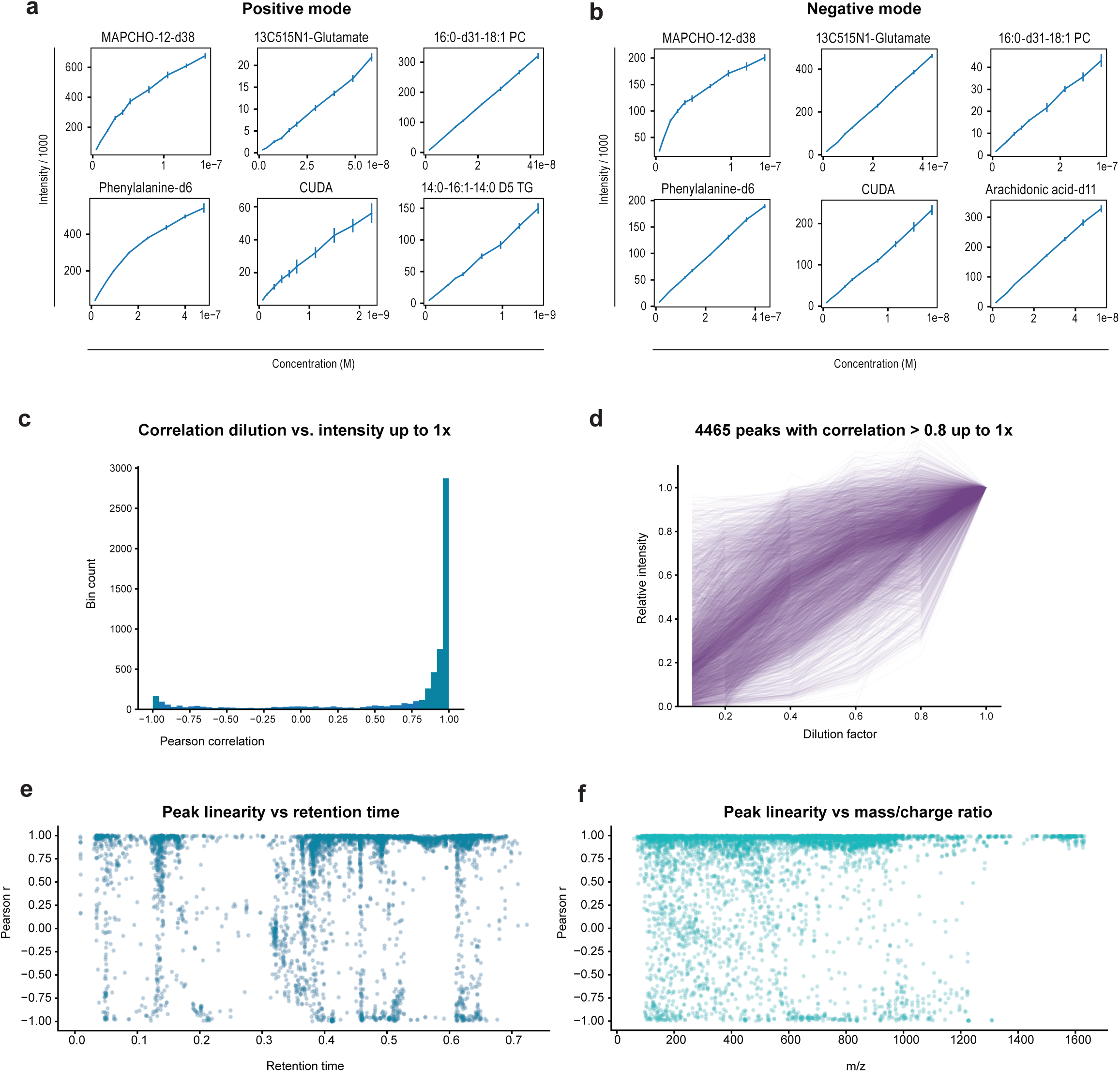
**a-b**, Dilution curves for internal standards in positive (a) and negative (b) ionization mode. Error bars denote standard deviation over n = 10 replicates. **c**, Linearity expressed as Pearson correlation with dilution factor for 6,178 rLC-MS features detected in human plasma. **d**, Dilution curves for 4,465 features with correlation > 0.8 in (a). **e**, Pearson correlation as in (a) vs. feature retention time. **f**, Pearson correlation as in (a) vs. mass/charge ratio (m/z).

**Supplementary Figure 2.**
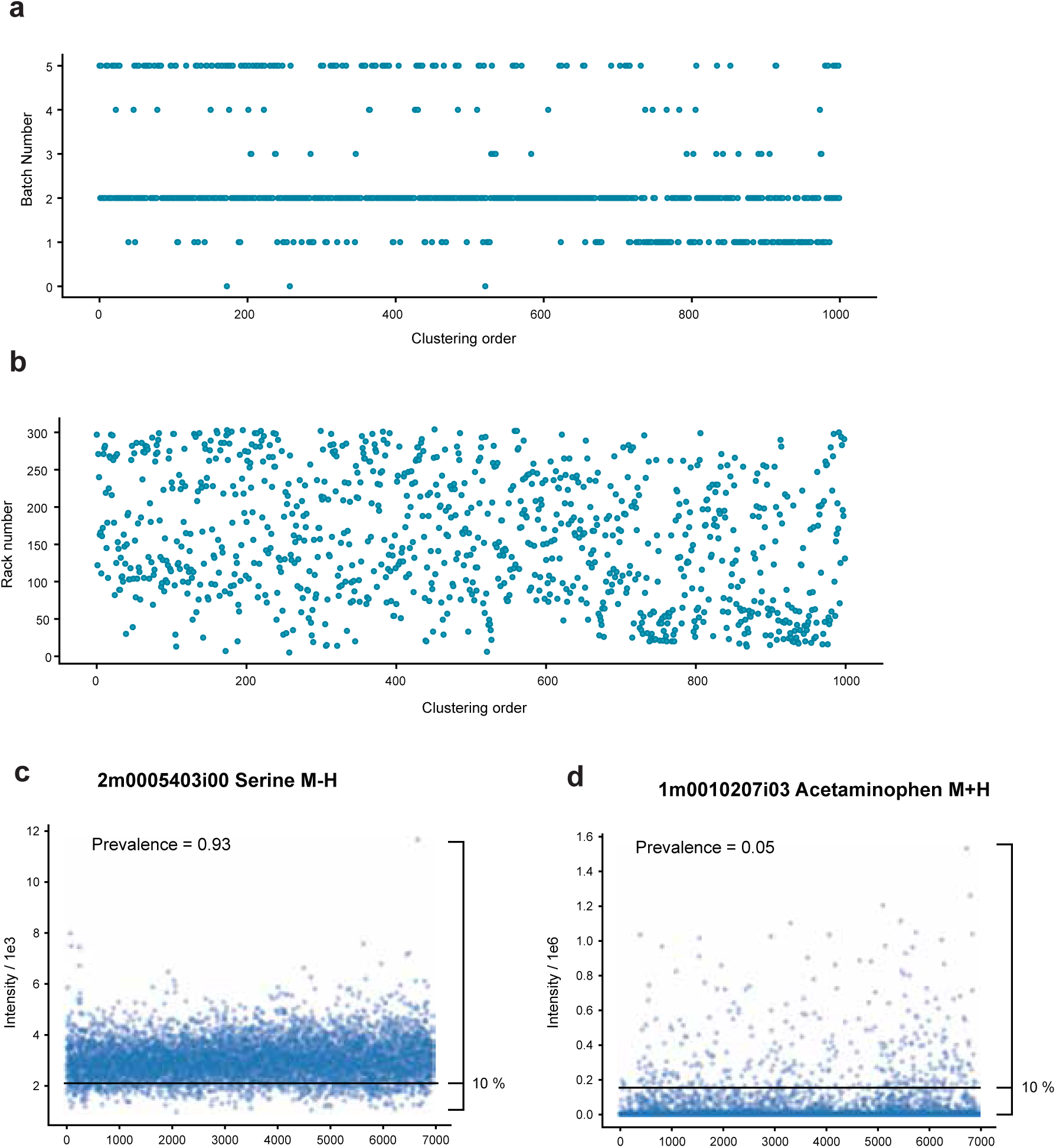
**a-b**, Batch number (a) and plate number (b) for a random subset of 1,000 samples from the human cohort, ordered by unsupervised hierarchical clustering. **c-d**, Examples feature intensity across individuals (mean over samples for each individual) for serine (c) and acetaminophen (d). Feature prevalence is defined as the fraction of individuals with feature values above a detection limit defined as 10% of the min-max range.

**Supplementary Figure 3.**
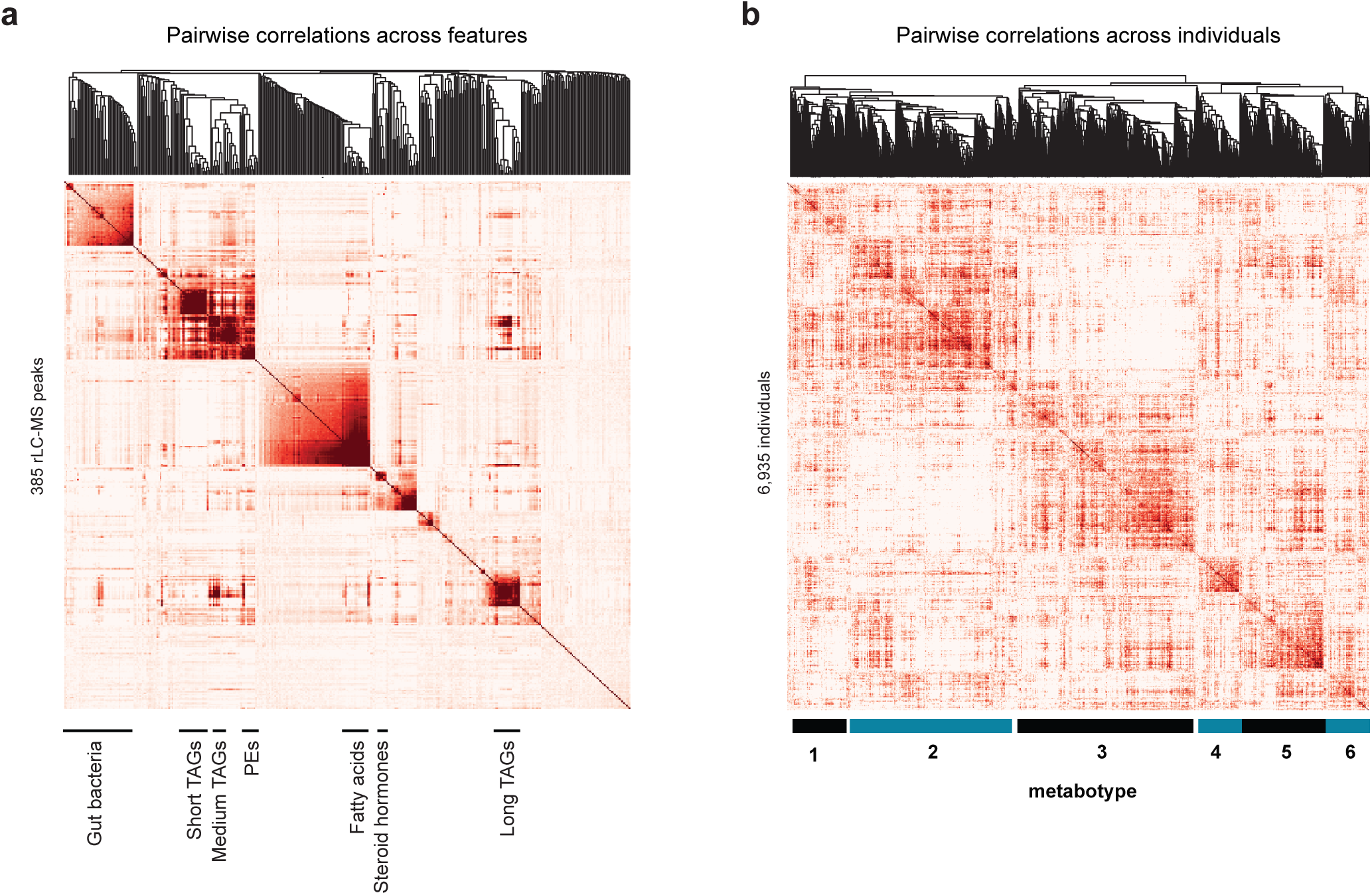
**a**, Clustered peak correlation matrix with clustering dendrogram. Metabolite clusters as in Fig 3 are indicated. **b**, Clustered individual correlation matrix with dendrogram. Metabotypes 1 to 6 as in Fig 3 are indicated.

**Supplementary Figure 4.**
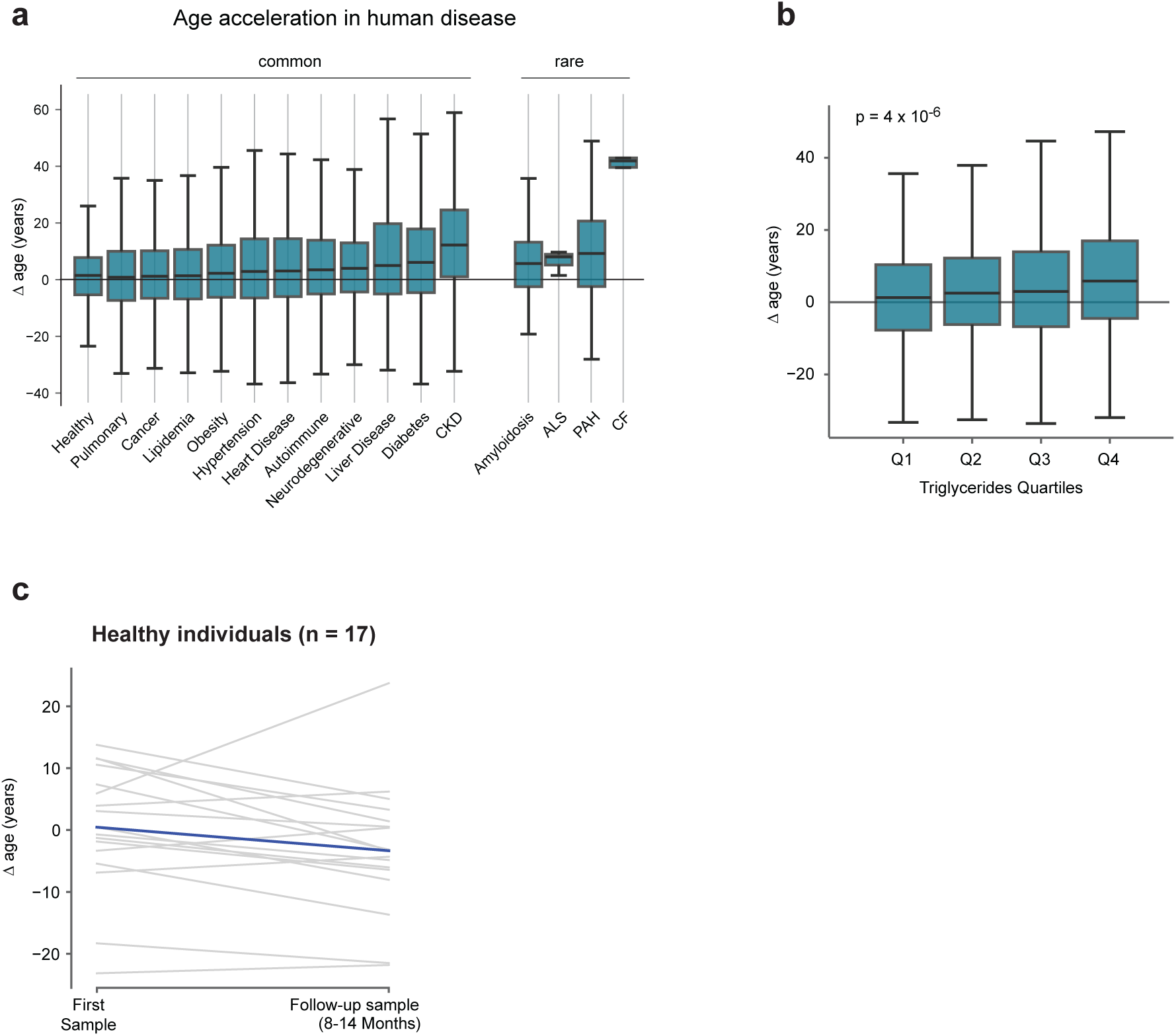
**a**, Distribution of age acceleration (Δage) for individuals diagnosed with indicated disorders, as in Fig 4c. Box plot indicating median, quartiles and min-max range are shown. **b**, Age acceleration *vs*. quartiles of plasma triglyceride level for individuals diagnosed with lipidemia. **c**, Age acceleration in healthy individuals over time.

## References

1. Wishart, D. S. Emerging applications of metabolomics in drug discovery and precision medicine. Nat. Rev. Drug Discov. 15, 473–484 (2016).

2. Nicholson, J. K. et al. Metabolic phenotyping in clinical and surgical environments. Nature 491, 384– 392 (2012).

3. Chen, L. et al. Metabolite discovery through global annotation of untargeted metabolomics data. Nat. Methods 18, 1377–1385 (2021).

4. da Silva, R. R., Dorrestein, P. C. & Quinn, R. A. Illuminating the dark matter in metabolomics. Proc. Natl. Acad. Sci. 112, 12549–12550 (2015).

5. Buergel, T. et al. Metabolomic profiles predict individual multidisease outcomes. Nat. Med. 28, 2309– 2320 (2022).

6. Wang, T. J. et al. Metabolite profiles and the risk of developing diabetes. Nat. Med. 17, 448–453 (2011).

7. Fuhrer, T., Heer, D., Begemann, B. & Zamboni, N. High-throughput, accurate mass metabolome profiling of cellular extracts by flow injection-time-of-flight mass spectrometry. Anal. Chem. 83, 7074– 7080 (2011).

8. Sarvin, B. et al. Fast and sensitive flow-injection mass spectrometry metabolomics by analyzing sample-specific ion distributions. Nat. Commun. 11, 3186 (2020).

9. Wishart, D. S. et al. HMDB 4.0: the human metabolome database for 2018. Nucleic Acids Res. 46, D608–D617 (2018).

10. Kantz, E. D., Tiwari, S., Watrous, J. D., Cheng, S. & Jain, M. Deep Neural Networks for Classification of LC-MS Spectral Peaks. Anal. Chem. 91, 12407–12413 (2019).

11. Assfalg, M. et al. Evidence of different metabolic phenotypes in humans. Proc. Natl. Acad. Sci. 105, 1420–1424 (2008).

12. Gavaghan, C. L., Holmes, E., Lenz, E., Wilson, I. D. & Nicholson, J. K. An NMR-based metabonomic approach to investigate the biochemical consequences of genetic strain differences: application to the C57BL10J and Alpk:ApfCD mouse. FEBS Lett. 484, 169–174 (2000).

13. Horvath, S. DNA methylation age of human tissues and cell types. Genome Biol. 14, 3156 (2013).

14. Argentieri, M. A. et al. Proteomic aging clock predicts mortality and risk of common age-related diseases in diverse populations. Nat. Med. 30, 2450–2460 (2024).

15. Lassen, J. K. et al. Large-Scale metabolomics: Predicting biological age using 10,133 routine untargeted LC–MS measurements. Aging Cell 22, e13813 (2023).

16. Lau, C.-H. E. et al. NMR metabolomic modeling of age and lifespan: A multicohort analysis. Aging Cell 23, e14164 (2024).

17. Wang, F. et al. Plasma metabolomic profiles associated with mortality and longevity in a prospective analysis of 13,512 individuals. Nat. Commun. 14, 5744 (2023).

18. Hédou, J. et al. Discovery of sparse, reliable omic biomarkers with Stabl. Nat. Biotechnol. 42, 1581–1593 (2024).

19. Xiao, H. et al. A Quantitative Tissue-Specific Landscape of Protein Redox Regulation during Aging. Cell 180, 968–983.e24 (2020).

20. Yang, H. et al. Gut microbial-derived phenylacetylglutamine accelerates host cellular senescence. *Nat*. Aging 5, 401–418 (2025).

21. Watrous, J. D. et al. Visualization, Quantification, and Alignment of Spectral Drift in Population Scale Untargeted Metabolomics Data. Anal. Chem. 89, 1399–1404 (2017).

